# Photorespiration pathways in a chemolithoautotroph

**DOI:** 10.1101/2020.05.08.083683

**Authors:** Nico J. Claassens, Giovanni Scarinci, Axel Fischer, Avi I. Flamholz, William Newell, Stefan Frielingsdorf, Oliver Lenz, Arren Bar-Even

## Abstract

Carbon fixation via the Calvin cycle is constrained by the side activity of Rubisco with dioxygen, generating 2-phosphoglycolate. The metabolic recycling of 2-phosphoglycolate, an essential process termed photorespiration, was extensively studied in photoautotrophic organisms, including plants, algae, and cyanobacteria, but remains uncharacterized in chemolithoautotrophic bacteria. Here, we study photorespiration in the model chemolithoautotroph *Cupriavidus necator* (*Ralstonia eutropha*) by characterizing the proxy-process of glycolate metabolism, performing comparative transcriptomics of autotrophic growth under low and high CO_2_ concentrations, and testing autotrophic growth phenotypes of gene deletion strains at ambient CO_2_. We find that the canonical plant-like C_2_ cycle does not operate in this bacterium and instead the bacterial-like glycerate pathway is the main photorespiratory pathway. Upon disruption of the glycerate pathway, we find that an oxidative pathway, which we term the malate cycle, supports photorespiration. In this cycle, glyoxylate is condensed with acetyl-CoA to give malate, which undergoes two oxidative decarboxylation steps to regenerate acetyl-CoA. When both pathways are disrupted, autotrophic growth is abolished at ambient CO_2_. We present bioinformatic data suggesting that the malate cycle may support photorespiration in diverse chemolithoautotrophic bacteria. This study thus demonstrates a so-far unknown photorespiration pathway, highlighting important diversity in microbial carbon fixation metabolism.

## Introduction

The Calvin cycle is responsible for the vast majority of carbon fixation in the biosphere. However, its activity is constrained by the low rate and the limited substrate specificity of its carboxylating enzyme Rubisco. The oxygenase side-activity of Rubisco converts ribulose 1,5-bisphosphate (RuBP) into 3-phosphoglycerate (3PG) and 2-phosphoglycolate (2PG), a dead-end metabolite which can inhibit the Calvin cycle (1, 2). The metabolic recycling of 2PG, termed photorespiration, is an essential process for most organisms that grow autotrophically via the Calvin cycle (3). Photorespiration has been extensively studied in photosynthetic organisms, including plants, algae, and cyanobacteria (4–7). The only identified photorespiration pathway in plants is the so-called C_2_ cycle (Fig. 1), in which 2PG is first dephosphorylated to glycolate, then oxidized to glyoxylate, and subsequently aminated to glycine. One glycine molecule is decarboxylated to give 5,10-methylene-THF which reacts with another glycine to yield serine. Serine is then deaminated to hydroxypyruvate, further reduced to glycerate, and finally phosphorylated to generate the Calvin cycle intermediate 3PG.

**Figure 1.**
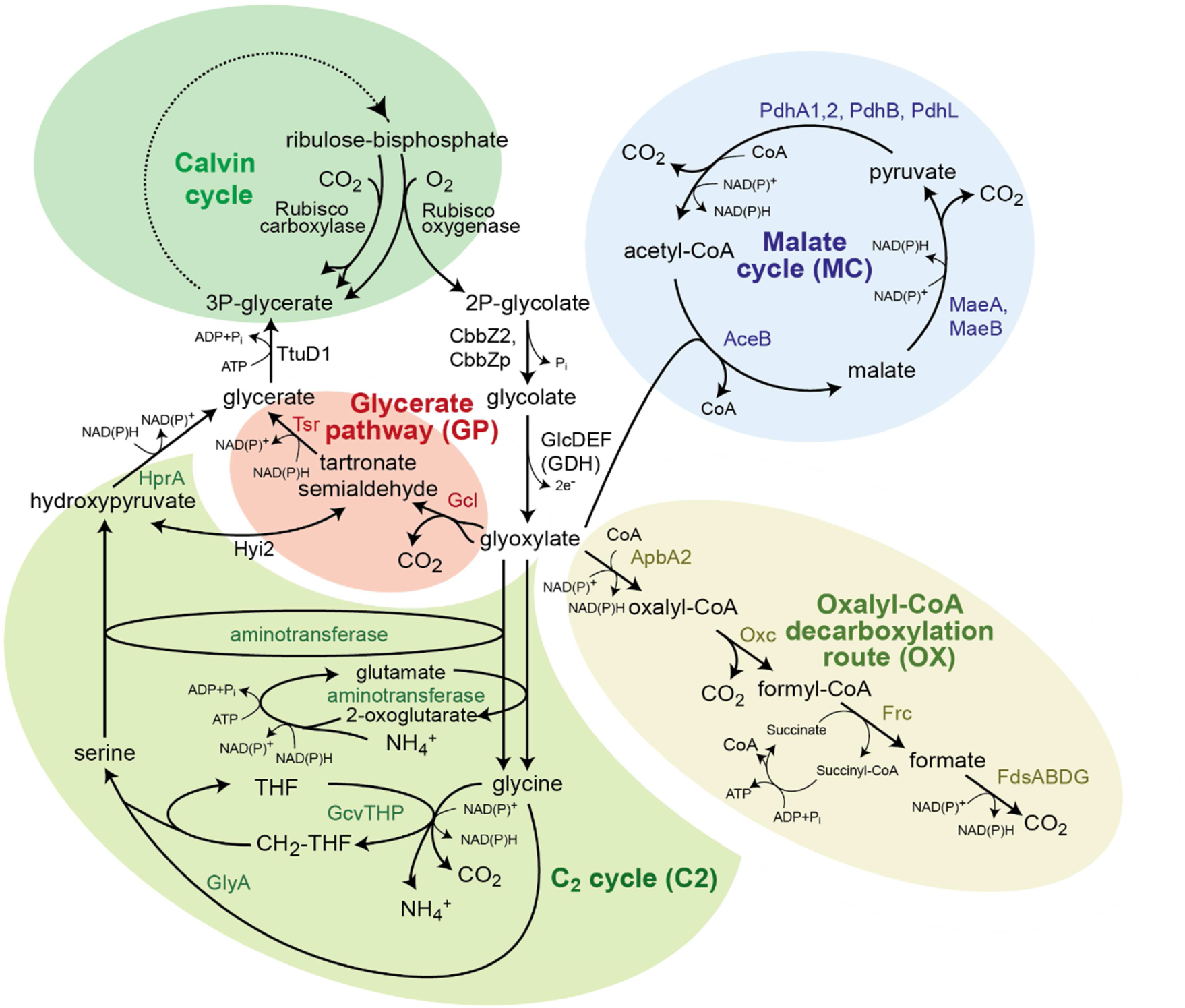
Candidate pathways supporting photorespiration in *C. necator*. 2-phosphoglycolate is first dephosphorylated to glycolate and then oxidized, by the glycolate dehydrogenase complex, to give glyoxylate. Glyoxylate can be further metabolized via four routes: the C_2_ cycle (i.e., the plant photorespiration pathway), the glycerate pathway, the oxalyl-CoA decarboxylation pathway, and malate cycle. The latter two routes completely oxidize glycoxylate to CO_2_. Note that that malate cycle may also proceed via oxidation of malate to oxaloacetate, which can be converted to pyruvate either directly or via phosphoenolpyruvate. Abbreviated enzyme names: AceB, malate synthase; ApbA2, CoA-acylating glyoxylate dehydrogenase; CbbZ2, CbbZp, 2-phosphoglycolate phosphatase; FdsABDG, formate dehydrogenase complex; Frc, formyl-CoA transferase; Gcl, glyoxylate carboligase; GcvTHP, glycine cleavage systems; GlcDEF, glycolate dehydrogenase complex; GlyA, serine hydroxymethyltransferase; HprA, hydroxypyruvate reductase; Hyi2, hydroxypyruvate isomerase; MaeA, MaeB, malic enzyme A/B; Oxc, oxalyl-CoA decarboxylase; PdhA1,2, PdhB, PdhL,, pyruvate dehydrogenase complex; Rubisco, Ribulose-bisphosphate carboxylase/oxygenase; Tsr, tartronate semialdehyde reductase; and TtuD1, glycerate kinase.

Using gene deletion studies, the cyanobacterium *Synechocystis sp.* PCC6803 was demonstrated to harbor two photorespiratory routes in addition to the C_2_ cycle (5, 8). In the glycerate pathway, two glyoxylate molecules are condensed to tartronate semialdehyde, which is subsequently reduced to glycerate and phosphorylated to 3PG (Fig. 1). Alternatively, in the oxalate decarboxylation pathway, glyoxylate is oxidized to oxalate and decarboxylated to formate, which is finally oxidized to CO_2_ (Fig. 1). Growth at ambient CO_2_ was abolished only when all three pathways were deleted, indicating that each of these routes can participate in photorespiration (5).

Photorespiration was initially named to describe light-dependent CO_2_ release in plants (9). Even though the name implies otherwise, photorespiration is not restricted to phototrophs but must occur in all organisms that use the Calvin cycle for autotrophic growth in the presence of oxygen. In fact, a wide range of chemolithoautotrophic microorganisms that employ the Calvin cycle under oxic conditions – including bacteria that oxidize hydrogen, ferrous iron, sulfur, or ammonia (10, 11) – must also cope with the oxygenase side-activity of Rubisco by recycling or removing 2PG. Yet, despite the obvious physiological significance of photorespiration to chemolithoautotrophs, it has so far received only scarce attention.

In this study, we explore metabolic routes involved in photorespiration of *Cupriavidus necator* H16 (formerly known as *Ralstonia eutropha* or *Alcaligenes eutrophus*), the best-studied chemolithoautotrophic microorganism that fixes CO_2_ fixation via the Calvin cycle (12–14). Unlike cyanobacteria, *C. necator* does not harbor a CO_2_ concentrating mechanism (i.e., a carboxysome with appropriate inorganic carbon transporters), as evident from the relatively high CO_2_ specificity of its Rubisco, which falls within the range reported for plants but is much higher than that found in cyanobacteria (15–17). Very little is known about photorespiration in *C. necator*. Previous studies found evidence only for the first steps of photorespiration – 2PG dephosphorylation and glycolate oxidation (18–20). However, the specific routes that metabolize glyoxylate remain elusive.

Here, we identify the native photorespiration pathways of *C. necator* by performing comparative transcriptomic analysis and conducting growth experiments with gene deletion strains. We show that the C_2_ cycle and decarboxylation via oxalate do not support photorespiration in *C. necator*. We find that the glycerate pathway is the primary photorespiration route in this bacterium. A second pathway, which we term the malate cycle, carries all photorespiratory flux when the glycerate pathway is disrupted. This route was previously unknown to operate in nature and can completely oxidize glyoxylate to CO_2_. In this cycle, glyoxylate is condensed with acetyl-CoA to generate malate, which then undergoes oxidative decarboxylation twice to regenerate acetyl-CoA (Fig. 1). Only when both the glycerate pathway and the malate cycle are disrupted is autotrophic growth at ambient CO_2_ abolished. This study therefore fills an important gap in our understanding of chemolithoautotrophic metabolism.

## Results

### Multiple pathways in *C. necator* support growth on glycolate

As glycolate metabolism is at the core of photorespiration (Fig. 1), we decided to start by exploring the metabolic pathways that can support the growth of *C. necator* on this C_2_ carbon source. First, we focused on the initial oxidation of glycolate to glyoxylate, which is the first step in all glycolate-metabolizing routes. The operon *glcDEF* is annotated to encode different subunits of the glycolate dehydrogenase complex (21). The gene *kch*, which is also part of the operon, is annotated as an ion transport channel, but as it shows homology to FAD-binding proteins, including subunit *D* of glycolate dehydrogenase from *Ralstonia syzygii* R24, it is more likely to encode another subunit of the complex. While a wild-type *C. necator* could efficiently grow on glycolate (doubling time of 3.2 ± 0.1 h, ‘WT’ in Fig. 2a), a strain deleted in the *glcDEF* operon failed to grow on this carbon source (‘∆GDH’ in Fig. 2a). This confirms that GlcDEF plays an essential role in oxidizing glycolate to glyoxylate.

**Figure 2.**
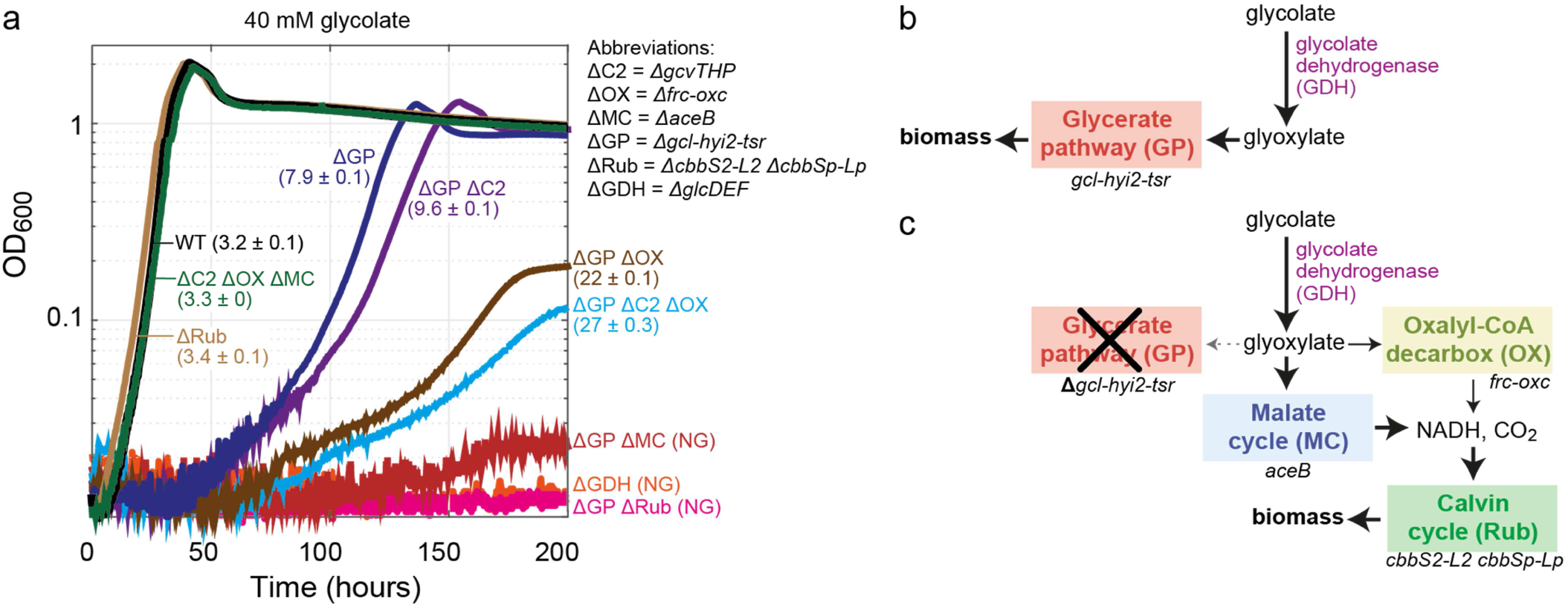
Growth of *C. necator* gene-deletion strains on glycolate. **(a)** Growth experiments were conducted in 96-well plate readers in minimal medium (JMM) supplemented with 40 mM sodium glycolate. Doubling times (hours) and standard deviations of biological triplicates are shown in between brackets, ‘NG’ corresponds to ‘no growth’. Triplicate growth experiments showed identical growth curves (±5%); hence, representative curves are shown. **(b)** The main route glycolate assimilation in *C. necator* growth is glycolate dehydrogenase followed by the glycerate pathway. **(c)** In the absence of the glycerate pathway, glyoxylate is decarboxylated via the oxalyl-CoA decarboxylation pathway and the malate cycle, the latter route being the dominant one. Generated CO_2_ and reducing equivalents are utilized by the Calvin cycle to support biomass formation. Strain labels: ∆C2, C_2_ cycle knockout (∆*gcvTHP*); ∆GDH, glycolate dehydrogenase knockout (∆*glcD-kch-glcE-glcF*); ∆GP, glycerate pathway knockout (∆*gcl-hyi-tsr*); *∆*MC, malate cycle knockout (∆*aceB*); *∆*OX, oxalate decarboxylation knockout (*frc-oxc*); *∆*Rub, Rubisco knockout (∆*cbbS2-cbbL2 ∆cbbSp-∆cbbLp*); WT, wild-type.

Next, we explored possible routes for glyoxylate metabolism. *C. necator* harbors genes encoding all the enzymes of the glycerate pathway in one operon: *gcl* (encoding glyoxylate carboligase), *hyi2* (hydroxypyruvate isomerase), *tsr* (tartronate semialdehyde reductase), and *ttuD1* (glycerate kinase). *C. necator* also harbors the key components of the C_2_ cycle, that is, the glycine cleavage system (encoded by *gcvTHP*) and serine hydroxymethyltransferase (*glyA*). While no dedicated glycine-dependent or serine-dependent transaminases are annotated in the genome of *C. necator* (Fig. 1), it is possible that glyoxylate amination and serine deamination are supported by one or more of the endogenous transaminase enzymes.

Finally, the *Synechocystis*-like oxalate decarboxylation pathway seems not to be present in *C. necator*, as no oxalate decarboxylase gene could be found. However, a similar route might be used. Specifically, oxalyl-CoA decarboxylase and formyl-CoA transferase – the genes for which, *oxc* and *frc*, are annotated and found in a single operon – could enable a glyoxylate oxidation route. In this putative route, oxalate is activated to oxalyl-CoA, which is then decarboxylated to formyl-CoA, converted to formate, and finally oxidized to CO_2_ (Fig. 1). However, no CoA-acylating glyoxylate dehydrogenase (catalyzing the first reaction of the pathway) is annotated in the genome of *C. necator*. Hence, we performed a BLAST search of the bacterium’s genome using a CoA-acylating glyoxylate dehydrogenase from *Methylobacterium extorquens* (*panE2)* (22) as a query. We identified the gene *apbA2*, located in close proximity to the *frc-oxc* operon and showing high sequence homology to the *M. extorquens* gene (66% similarity, 51% identity), as a probable candidate glyoxylate dehydrogenase.

To test whether ApbA2 can indeed catalyze the reversible CoA-acylating glyoxylate dehydrogenase reaction *in vivo*, we tested the growth of a *∆apbA2* strain on oxalate. Growth on oxalate can proceed via two routes (Supplementary Fig. S1): (i) assimilation to central metabolism via the glycerate pathway, which depends on the activity of a CoA-acylating glyoxylate dehydrogenase (in the reverse direction of that required for growth on glycolate); and (ii) complete oxidation of oxalate which can be followed by carbon fixation via the Calvin cycle. Hence, if ApbA2 serves as a CoA-acylating glyoxylate dehydrogenase, its deletion should impair growth on oxalate. Indeed, while a wild-type *C. necator* grew efficiently on oxalate (doubling time 6.5 ± 0.1, Supplementary Fig. S1) growth of the ∆*apbA2* strain on this carbon source was impaired (doubling time of 10 ± 1.1 h, Supplementary Fig. S1). This deletion strain showed a similar growth phenotype to that observed upon deletion of the enzymes of the glycerate pathway (doubling time of 13 ± 0.7 h, Supplementary Fig. S1). It therefore seems that ApbA2 can indeed act as a CoA-acylating glyoxylate dehydrogenase. Interestingly, deletion of the *frc*-*oxc* operon completely abolished growth on oxalate (Supplementary Fig. S1), presumably as the supply of reducing power via oxalate oxidation is necessary to support the assimilation of this highly oxidized substrate.

By generating three distinct mutant strains, carrying deletions in the *gcl*-*hyi2*-*tsr* operon, the *gcvTHP* operon, or the *frc-oxc* operon, we explored the contribution of each candidate route to growth on glycolate. While the deletion of the latter two operons did not affect growth on glycolate (doubling time of 3.4 ± 0.1 h, ‘∆C2 ∆OX ∆MC’ in Fig. 2a), deletion of the *gcl*-*hyi2*-*tsr* operon resulted in a substantially lower growth rate (doubling time of 7.9 ± 0.1 h, ‘∆GP’ in Fig. 2a), indicating that the glycerate pathway is the main route for growth on glycolate (Fig. 2b). Still, the ability of the strain lacking the *gcl*-*hyi2*-*tsr* operon to grow on glycolate implies that other routes can support glyoxylate metabolism.

Next, we deleted the *gcvTHP* operon or the *frc-oxc* operon in the strain already lacking the *gcl*-*hyi2*-*tsr* operon. We found that further disruption of the oxalyl-CoA decarboxylation pathway substantially reduced the growth rate (doubling time of 22 ± 0.1 h, ‘∆GP ∆OX’ in Fig. 2a), indicating that this route supports glyoxylate metabolism in the absence of the glycerate pathway. On the other hand, deletion of *gcvTHP*, disrupting the C_2_ pathway, had only a small negative effect on growth (doubling time of 9.6 ± 0.1 h, ‘∆GP ∆C2’ in Fig. 2a), suggesting that this route contributes only marginally to the metabolism of glyoxylate.

Even after deleting all three operons, growth on glycolate was still observed (doubling time of 27 ± 0.1 h, ‘∆GP ∆C2 ∆OX’ in Fig. 2a), indicating that additional unknown route(s) can support glyoxylate metabolism. Yet, only few other enzymes can react with glyoxylate and even fewer exist in *C. necator*. For example, the β-hydroxyaspartate cycle was recently shown to enable growth on glycolate via glyoxylate assimilation (23), but the genome of *C. necator* does not encode its key glyoxylate assimilating enzyme, β-hydroxyaspartate aldolase. We were able to identify only one other enzyme in this bacterium that can react with glyoxylate: malate synthase, a key component of the glyoxylate shunt that condenses glyoxylate with acetyl-CoA to generate malate (24, 25). Indeed, when the gene encoding malate synthase (*aceB*) was deleted in the strain deleted in the *gcl*-*hyi2*-*tsr* operon, growth on glycolate was completely abolished (‘∆GP ∆MC’ in Fig. 2a). On the other hand, deletion of *aceB* in a strain in which the glycerate pathway is still active did not affect growth on glycolate (‘∆C2 ∆OX ∆MC’ in Fig. 2a).

For glyoxylate metabolism via malate synthase to proceed, the co-substrate acetyl-CoA must be regenerated. Such regeneration requires malate to undergo oxidative decarboxylation twice, first to pyruvate and then to acetyl-CoA. The combined activity of the malic enzyme and pyruvate dehydrogenase can support this double oxidation. The result is a ‘malate cycle’, composed of malate synthase, malic enzyme, and pyruvate dehydrogenase, which together completely oxidize glyoxylate to CO_2_ while generating two NAD(P)H molecules (Fig. 1). (We note that this cycle could alternatively proceed via malate oxidation to oxaloacetate, which is then converted to phosphoenolpyruvate via phosphoenolpyruvate carboxykinase, and further metabolized to pyruvate and acetyl-CoA; this alternative malate cycle would result in the same net decarboxylation reaction).

*C. necator* is expected to grow on glycolate via the malate cycle only by using the generated reducing power to fix CO_2_ via the Calvin cycle (Fig. 2c). To test if this is indeed the case, we deleted all genes encoding for Rubisco (*cbbS2*, *cbbL2*, *cbbSp*, and *cbbLp*) in the strain disrupted in the glycerate pathway (i.e., deleted in the *gcl*-*hyi2*-*tsr* operon). As anticipated, this strain was not able to grow on glycolate (‘∆GP ∆Rub’ in Fig. 2a); note that the deletion of Rubisco in a wild-type strain did not affect growth on glycolate as the glycerate pathway is still active (doubling time of 3.3 h, ‘∆Rub’ in Fig. 2a). These results confirm that growth on glycolate via the malate cycle strictly depends on the carbon fixation via the Calvin cycle.

To summarize, growth on glycolate mainly depends on the glycerate pathway in a wild-type *C. necator* (Fig. 2a). When this route is disrupted, glyoxylate is metabolized via a combination of the malate cycle and the oxalyl-CoA decarboxylase pathway, both of which depend on CO_2_ fixation for growth (Fig. 2c). Under these conditions, the malate cycle carries most of the flux and is essential for growth, while the oxalyl-CoA decarboxylase pathway has a secondary role. Yet, it is still not clear whether the pathways that support growth on glycolate also participate in photorespiration during autotrophic growth at ambient CO_2_.

### Photorespiration in *C. necator* is supported by the glycerate pathway and the malate cycle

To study which of the pathways of glycolate metabolism also participates in photorespiration, we compared the transcript levels of a wild-type *C. necator* growing autotrophically on hydrogen at ambient CO_2_ concentrations (≈0.04%) *versus* elevated CO_2_ concentrations (10%); the latter condition suppresses the oxygenation reaction. We found that the genes encoding for the Calvin cycle enzymes were overexpressed between 4- and 12-fold under ambient CO_2_ concentration (Supplementary Data 1). This result is expected as higher activity of the cycle is needed to compensate for the decreased rate of Rubisco and the carbon loss from photorespiration. Furthermore, the genes encoding the first steps of photorespiration, that is, 2PG phosphatase (*cbbZ2* and *cbbZp*) and the glyoxylate dehydrogenase complex, were overexpressed between 4- and 8-fold (Fig. 3 and Supplementary Data 1).

**Figure 3.**
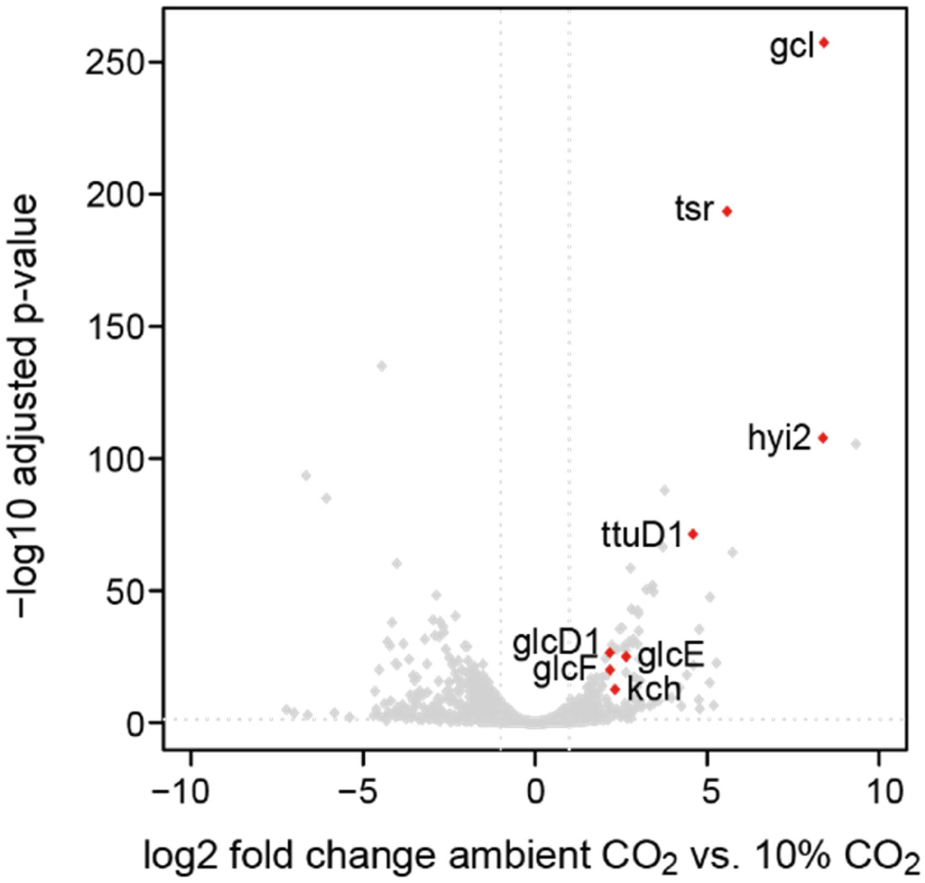
Volcano plot comparing the *C. necator* transcriptome in autotrophic growth at ambient CO_2_ versus 10% CO_2_. RNA was isolated during mid-log phase from *C. necator* grown in shake-flask cultures on minimal medium (JMM) with a headspace consisting of ambient air and 4% hydrogen or 10% CO_2_, 4% hydrogen. Differential expression analysis was performed as described in the materials and methods, log2-transformed fold changes (log2fc) and adjusted p-values of all genes are depicted in the graph. Only genes that are significantly upregulated and related to photorespiration pathways are labelled in the plot (red dots). Values for all genes can be found in Supplementary Data 1. Gene abbreviations and encoded enzymes: *gcl*, glyoxylate carboligase; *glcD1,glcE,glcF*, glycolate dehydrogenase complex subunits; *hyi2*, hydroxypyruvate isomerase; *kch*, putative glycolate dehydrogenase subunit; *oxc*, oxalyl-CoA decarboxylase; *tsr*, tartronate semialdehyde reductase; *ttuD1*, glycerate kinase.

Genes encoding the enzymes of the glycerate pathway were among the most highly upregulated at ambient CO_2_ (Fig. 3 and Supplementary Data 1): *gcl* and *hyi2* were more than 300-fold overexpressed, *tsr* was ≈50-fold upregulated, and *ttuD1* was ≈25-fold upregulated. On the other hand, the genes related to other potential glyoxylate metabolism routes – that is, the C_2_ pathway, the oxalyl-CoA decarboxylation pathway, and the malate cycle – were not overexpressed at ambient CO_2_ (Supplementary Data 1). This suggests that the glycerate pathway is the main photorespiration route. However, the fact that the genes of the other pathways were not overexpressed does not necessarily mean that they do not participate in photorespiration. Specifically, it could be that their basal expression levels are sufficient to support the required activity. For example, the genes encoding for the components of the malate cycle – *aceB*, *maeA*, *maeB*, *pdhA1*, *pdhB*, and *pdhL –* are highly expressed both under ambient and high CO_2_ concentrations (all among the 10% most highly expressed in both conditions; see Supplementary Data 1); hence, the malate cycle could play a role in photorespiration.

To determine the relative importance of each candidate pathway in photorespiration, we tested wild-type *C. necator* and several of the gene deletion strains described above for their ability to grow autotrophically at ambient CO_2_ (Fig. 4). Autotrophic growth of wild-type *C. necator* under these conditions resulted in a much lower growth rate than observed at 10% CO_2_ (doubling time of 21 ± 0.7 h *vs*. ~3 h, solid *vs*. dashed ‘WT’ lines in Fig. 4, respectively). This difference is expected from the lower carboxylation rate of Rubisco at low CO_2_ concentrations and the relatively high rate of the oxygenation reaction which leads to CO_2_ release, thus directly counteracting carbon fixation.

**Figure 4.**
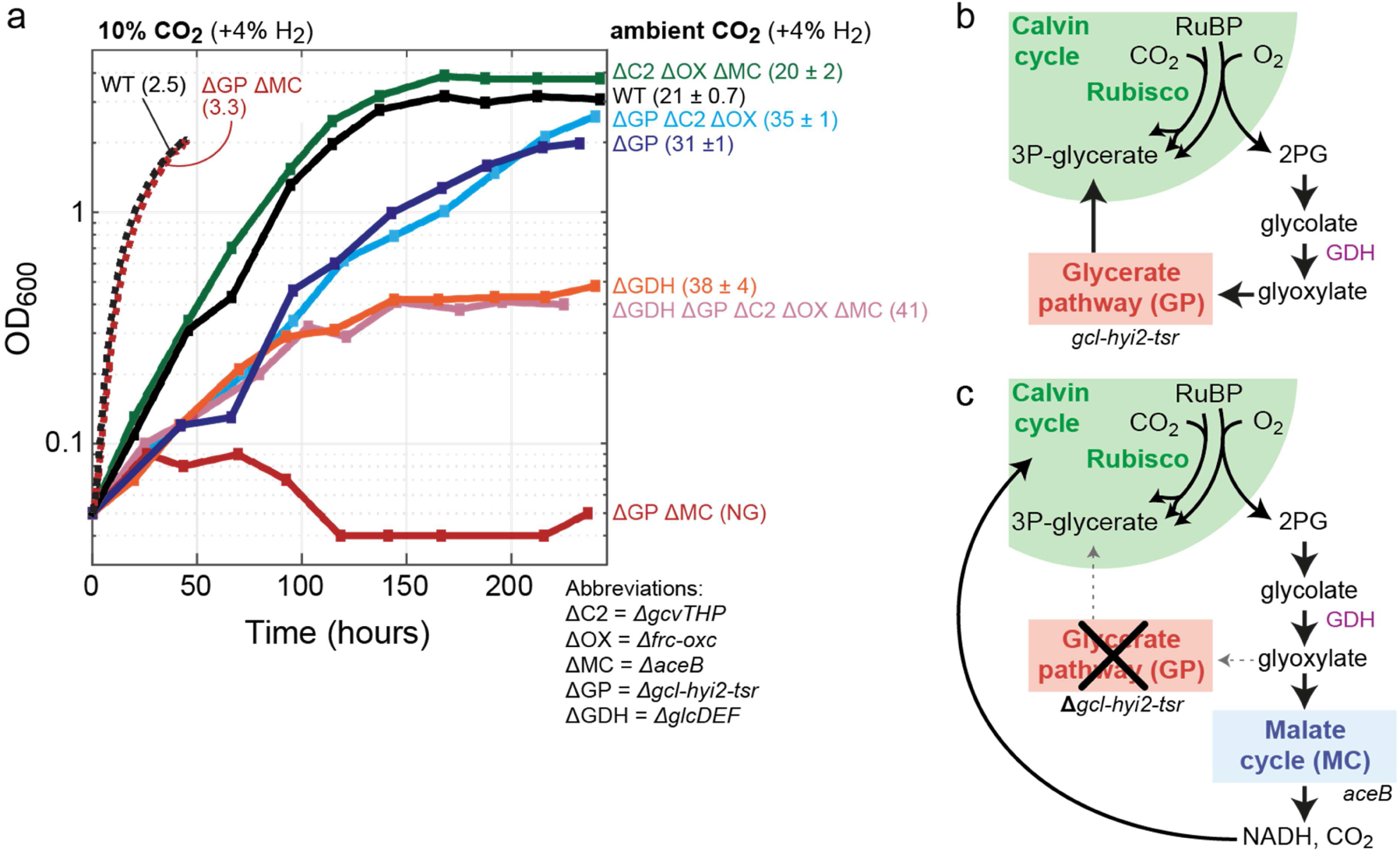
Autotrophic growth of *C. necator* gene-deletion strains under ambient CO_2_. **(a)** Growth experiments were conducted in 700 mL bioreactor cultures on minimal medium (JMM) with a continuous sparging of gas (6.25 L/minute) with ambient air + 4% hydrogen; control experiments were conducted at 10% CO_2_ + 4 % hydrogen (supplemented with air). Doubling times (hours) and standard deviations are shown in between brackets, ‘NG’ corresponds to ‘no growth’. Growth experiments on ambient CO_2_ were performed in biological duplicates and showed identical growth curves (±5%); hence, representative curves are shown. We performed control growth experiments on high CO_2_ for two strains only once and only up to 50 hours to limit extremely high CO_2_ turnover in the (non-recycling) bioreactor set-up. **(b)** Photorespiration in *C. necator* proceeds mainly via glycolate dehydrogenase and the glycerate pathway to regenerate 3-phosphoglycerate (3P-glycerate) for the Calvin cycle**. (c)** In the absence of the glycerate pathway, glyoxylate is decarboxylated by the malate cycle to CO_2_ and NADH, which can be reassimilated by the Calvin cycle. Strain labels: ∆C2, C_2_ cycle knockout (∆*gcvTHP*); ∆GDH, glycolate dehydrogenase knockout (∆*glcD-kch-glcE-glcF*); ∆GP, glycerate pathway knockout (∆*gcl-hyi-tsr*); ∆MC, malate cycle knockout (∆*aceB*); ∆OX, oxalate decarboxylation knockout (∆*frc-oxc*); WT, wild-type.

A strain lacking all routes of glyoxylate metabolism besides the glycerate pathway did not show reduced growth at ambient CO_2_ (doubling time of 20 ± 2 h, ‘∆C2 ∆OX ∆MC’ in Fig. 4). On the other hand, a strain in which the glycerate pathway was disrupted displayed a substantially lower growth rate (doubling time of 35 ± 1 h, ‘∆GP’ in Fig. 4). This suggests that the glycerate pathway is the major route of photorespiration, but also that it can be replaced by other pathways. Further deletion of the malate cycle in the strain lacking the glycerate pathway completely abolished autotrophic growth at ambient CO_2_ (‘∆GP ∆MC’ in Fig. 4). On the other hand, disruption of the C_2_ pathway and the oxalyl-CoA decarboxylation pathway in the strain lacking the glycerate pathway did not alter its growth phenotype (doubling time of 31 ± 1 h, ‘∆GP ∆C2 ∆OX’ in Fig. 4). These results clearly show that the malate cycle can participate in photorespiration while the other two pathways contribute little, if any, to this process. Notably, deletion of both the glycerate pathway and the malate cycle did not affect autotrophic growth at high CO_2_ concentrations (dashed ‘∆GP ∆MC’ line in Fig. 4), as photorespiration is expected to be negligible under these conditions due to competitive inhibition of oxygenation at high CO_2_.

We were also interested to explore the outcome of disrupting photorespiratory metabolism upstream of glyoxylate. We found that a strain deleted in glycolate dehydrogenase could grow autotrophically under ambient CO_2_ concentration, albeit at substantially lower growth rate and yield (doubling time of 38 ± 4 h, ‘∆GDH’ in Fig. 4). Supporting previous studies (18, 19), we found that glycolate accumulates in the medium during autotrophic growth of this strain at ambient CO_2_ (Fig. 5). Since glycolate is secreted by this strain and is not further oxidized, we hypothesized that further deletion of all four glyoxylate metabolism routes would not affect the growth phenotype; we indeed found this to be the case (doubling time of 41 h, ‘∆GDH ∆GP ∆C2 ∆OX ∆MC’ in Fig. 4). Moreover, as expected, no glycolate was detected during the autotrophic growth of a wild-type strain at ambient CO_2_.

**Figure 5.**
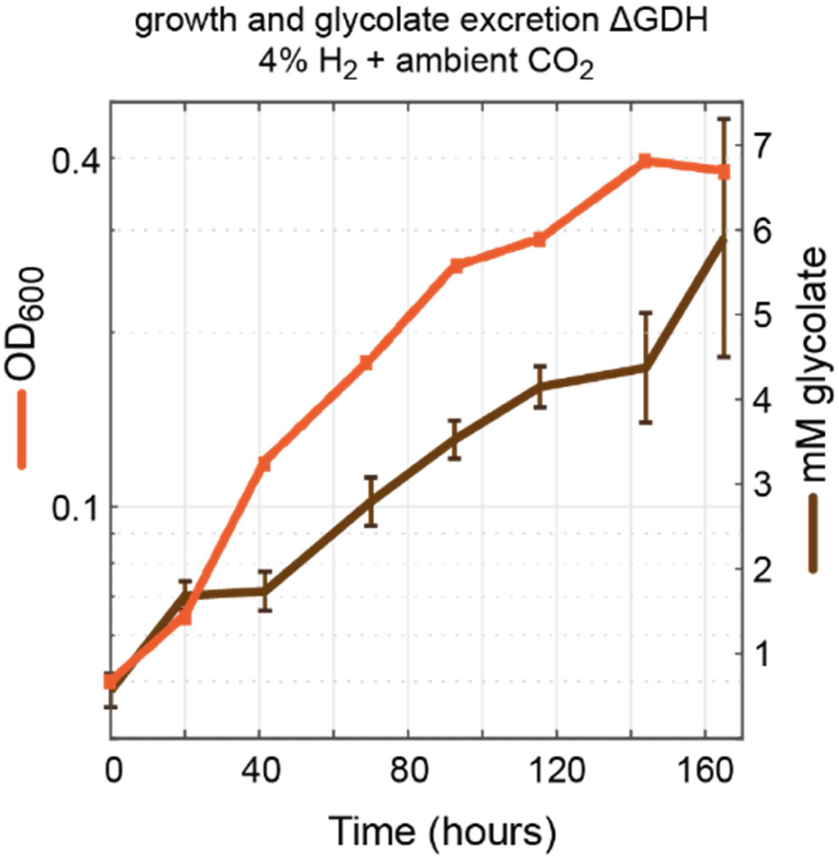
Glycolate excretion during autotrophic growth on the glycolate dehydrogenase knockout strain. Growth (OD_600_) and glycolate concentrations were monitored every 24 hours during the growth phase of a *C. necator* ∆*glcD-kch-glcE-glcF* strain. Growth experiments were conducted in 700 mL bioreactor cultures on minimal medium (JMM) with a continuous sparging of gas (6.25 L/minute) with ambient air + 4% hydrogen. Biological duplicates were measured and SD for the glycolate concentrations are shown, replicates showed identical growth curves (±5%) and a representative curve is shown. Glycolate was measured by ion chromatography as explained in materials and methods.

It therefore seems that *C. necator* can excrete glycolate if necessary, thus enabling the autotrophic growth of a strain lacking glycolate dehydrogenase. In contrast, *C. necator* seems to be incapable of secreting glyoxylate, which prevents the growth of a strain lacking the glycerate pathway and the malate cycle. Indeed, we could not detect glyoxylate in the growth medium of any tested strains.

### The malate cycle may be prevalent in chemolithoautotrophs using the Calvin cycle

To explore the potential distribution of the malate cycle, we searched for the occurrence of malate synthase (PFAM 01274 (26)) in bacteria that harbor a bona-fide Rubisco, and are therefore likely to operate the Calvin cycle and rely on a photorespiration to metabolize 2PG (Methods). Only 2% of cyanobacterial species harbor malate synthase, indicating that the malate cycle is likely uncommon among oxygenic photoautotrophic bacteria (Table 1). However, of the remaining ≈2000 non-cyanobacterial genomes found to encode Rubisco, ≈60% also encode a malate synthase. These include bacteria that grow chemolithoautotrophically by oxidizing either inorganic compounds (e.g., hydrogen, ammonia, and sulfur compounds) or organic one-carbon compounds (e.g., formate or methanol), as well as non-oxygenic phototrophs that use the Calvin cycle (such as purple non-sulfur bacteria). In several key orders of aerobic autotrophic bacteria, most genomes that harbor a Rubisco also encode a malate synthase (Table 1), including *Burkholderiales* (73%), *Chromatiales* (70%), *Rhizobiales* (98%), *Rhodobacterales* (88%), *Thiotrichales* (100%), *Mycobacteriales* (98%), and *Sulfobacilliales* (71%). While the presence of malate synthase does not necessarily indicate that the malate cycle operates, sufficient expression levels of this enzyme under relevant conditions would probably lead to a substantial photorespiratory flux via this route.

**Table 1.** Prevalence of malate synthase in chemolithoautotrophs and cyanobacteria that use the Calvin cycle. Shown are only orders that have many aerobic chemolithoautotrophs. Supplementary Data 2 shows all orders and species.

| Phylum | Order | Genera examples | Type of organisms | Electron donor | # with Rubisco | % with malate synthase |
| --- | --- | --- | --- | --- | --- | --- |
| **OXYGENIC PHOTOAUTOTROPHS** |  |  |  |  | **536** | **2%** |
| **Cyanobacteria** |  |  | photoautotroph | water (oxygenic photosynthesis) | 536 | 2% |
| **CHEMOLITHOAUTOTROPHS** |  |  |  |  | **1936** | **63%** |
| **Proteobacteria** | Acidithiobacillales | Acidithiobacillus | chemolithoautotrophs | iron, hydrogen, sulfur compounds | 9 | 0% |
|  | Burkholderiales | Cupriavidus, Nitrosomonas | chemolithoautotrophs, | hydrogen, formate, ammonia, sulfur compounds, iron | 405 | 73% |
|  | Chromatiales | Allochrocatium, Thiocapsa | chemolithoautotrophs, purple sulfur bacteria (anoxygenic phototrophs) | hydrogen, sulfur compounds | 40 | 70% |
|  | Ectothiorhodospirales | Thioalkalivibrio | chemolithoautotrophs, purple sulfur bacteria (anoxygenic phototrophs) | sulfur compounds | 35 | 40% |
|  | Halothiobacillales | Nitrosococcus | chemolithoautotrophs | sulfur compounds | 6 | 0% |
|  | Nitrosococcales | Halothiobacillus | chemolithoautotrophs | ammonia | 14 | 0% |
|  | Rhizobiales | Nitrobacter, Rhodopseudomonas | chemolithoautotrophs, purple non-sulfur bacteria (anoxygenic phototrophs), methylotrophs | hydrogen, nitrite, methanol, formate | 305 | 98% |
|  | Rhodobacterales | Rhodobacter, Paracoccus | chemolithoautotrophs, purple non-sulfur bacteria (anoxygenic phototrophs), methylotrophs | hydrogen, sulfur compounds, methanol | 150 | 88% |
|  | Rhodospirillales | Rhodospirillum | chemolithoautotrophs, purple non-sulfur bacteria (anoxygenic phototrophs) | hydrogen | 46 | 39% |
|  | Thiomicrospirales | Hydrogenvibrio | chemolithoautotrophs | hydrogen, sulfur compounds | 39 | 15% |
|  | Thiotrichales | Thiotrix | chemolithoautotrophs | sulfur compounds | 7 | 100% |
| **Actinobacteriota** | Mycobacteriales | Mycobacterium | methylotrophs, carboxidotrophs | carbon monooxide, methanol | 133 | 98% |
| **Firmicutes** | Sulfobacilliales | Sulfobacillus | chemolithoautotrophs | sulfur compounds | 7 | 71% |

The other enzymes of the malate cycle – the malic enzyme and pyruvate dehydrogenase or their alternatives (e.g., malate dehydrogenase, phosphoenolpyruvate carboxykinase) – are quite ubiquitous. However, these enzymes are not strictly necessary. While growth on glycolate via the malate cycle must be accompanied by the regeneration of acetyl-CoA and thus complete oxidation of glyoxylate, in photorespiration, acetyl-CoA does not need to be regenerated. Rather, the glyoxylate produced in photorespiration can be condensed with acetyl-CoA generated from the carbon fixation process and the resulting malate assimilated into biomass. In this case, the malate ‘cycle’ is not a real cycle but rather represents a linear photorespiration route to reassimilate glyoxylate into central metabolism. We suspect that some chemolithoautotrophs using malate synthase as part of their photorespiratory metabolism actually operate the linear than rather than cyclic version of this pathway.

## Discussion

This study aimed to fill gaps in our knowledge of photorespiratory metabolism in chemolithoautotrophic microorganisms. In the model chemolithoautotroph *C. necator*, we confirmed the role of the glycolate dehydrogenase complex in photorespiration and further revealed that two metabolic routes can sustain photorespiratory flux and thus support autotrophic growth at ambient CO_2_.

The glycolate dehydrogenase complex of *C. necator* is homologous to other bacterial glycolate dehydrogenases. The cofactor used as an electron acceptor for glycolate oxidation has not yet been elucidated in any bacteria. It is sometimes proposed, especially in cyanobacterial photorespiration, that NAD^+^ serves as the electron acceptor. However, this is highly doubtful as the change in Gibbs energy for the reaction glycolate + NAD^+^ = glyoxylate + NADH is very high (∆_r_G^’m^ > 40 kJ/mol, pH 7.5, ionic strength of 0.25 mM; ∆_r_G^’m^ corresponds to metabolite concentration of 1 mM (27)). Instead, it is more likely that glycolate transfers its electrons, via a flavin adenine dinucleotide cofactor, to a quinone, the reduction potential of which is substantially higher than that of NAD (E^’m^ ~ 0 mV rather than ~ −300 mV, respectively).

Unlike most phototrophic organisms, which mostly cannot grow heterotrophically, *C. necator* can be grown on various organic compounds. We used this metabolic versatility to explore photorespiration via a proxy-process of glycolate metabolism. Indeed, we found that photorespiration and growth on glycolate generally rely on the same routes, that is, the glycerate pathway and the malate cycle. In contrast, the oxalyl-CoA decarboxylation pathway seems to participate, albeit marginally, only in glycolate metabolism but not in photorespiration. This might be attributed to differences in regulation or to the higher concentrations of glyoxylate available when supplying glycolate as a sole carbon source.

Autotrophic growth phenotypes of our *C. necator* gene deletion strains show that the C_2_ pathway contributes negligibly to photorespiration and glycolate metabolism. While *C. necator* harbors the main components of this pathway, transaminase enzymes that accept glycine and serine could not be identified. It might be the case that these enzymatic activities are completely missing in the bacterium. Alternatively, it could be that endogenous transaminases can aminate glyoxylate and deaminate serine but the C_2_ pathway shows only low activity due to inadequate regulation.

A key finding of our work is the existence of the malate cycle as a supporting route for photorespiration and growth on glycolate. It is difficult to determine whether this route plays a role in the wild-type strain, as its deletion does not seem to hamper growth when the glycerate pathway is present. It could be that the malate cycle carries non-negligible flux only when the glycerate pathway is disrupted and glyoxylate begins to accumulate. Alternatively, considering the relative high expression of malate synthase, it is possible that the pathway always supports a substantial fraction of glyoxylate metabolism, but not enough to affect growth once deleted.

The glycerate pathway is the most efficient, naturally occurring photorespiration route in terms of consumption of ATP and reducing power (28). On the other hand, complete decarboxylation of glyoxylate to CO_2_ – as supported by the cyanobacterial oxalate decarboxylation pathway as well as the oxalyl-CoA decarboxylation pathway and the malate cycle described here – is arguably the least efficient photorespiration mode, as it requires higher activity of the Calvin cycle to compensate for the lost carbon. This might explain why the deletion of the glycerate pathway in *C. necator*, such that the malate cycle carries the entire photorespiratory flux, resulted in lower growth rate and yield (Fig. 4). Similarly, the superiority of the glycerate pathway might explain why it serves as the major photorespiration route both in cyanobacteria and in *C. necator*.

Despite the relative inefficiency of the malate cycle, its implementation in plants was suggested to boost carbon fixation and photosynthesis (28–31). Recently, the heterologous expression of malate synthase and glycolate dehydrogenase within the chloroplast of the agricultural crop *Nicotiana tabacum* led to a substantial increase in photosynthetic rate and yield (29). As the malate cycle is less efficient than the natural C_2_ pathway, this growth enhancement is not easy to explain and was suggested to be related to the release of CO_2_ in the chloroplast, rather than the mitochondria, thus suppressing the oxygenase side-reaction and promoting Rubisco’s carboxylation.

As malate synthase is present in many chemolithoautotrophs that use the Calvin cycle (Table 1), it is tempting to suggest that it contributes to photorespiration in many of these bacteria. However, the occurrence of malate synthase in cyanobacteria is much lower (Table 1). *Synechocystis*, the only bacterial photoautotroph for which photorespiration has been physiologically characterized (5, 8), probably lacks malate synthase and hence cannot operate the malate cycle (32, 33). However, the presence of malate synthase has been confirmed in some cyanobacteria (34–37), leading to the suggestion that they may use the malate cycle. We leave it for future studies to explore the occurrence of the malate cycle and other photorespiration routes in cyanobacteria and chemolithoautotrophs. Such investigations will generate further insights into the evolutionary history of photorespiratory metabolism.

## Materials and methods

### Strains, conjugations and gene deletions

*C. necator* H16 (DSMZ 428) was used for transcriptome studies. Growth experiments were performed for a *C. necator* H16 strain knocked out for polyhydroxybutyrate biosynthesis (∆*phaC1*) (38), in which other gene deletions were performed. The ∆*phaC1* strain grows in nutrient non-limiting similar to the wild-type and does not result in PHB granules that could disturb optical density measurements.

Cloning of plasmids was performed in *E. coli* DH5α, whereas *E. coli* S17-1 was used for conjugation of mobilizable plasmids to *C. necator* by biparental overnight spot mating. A complete overview of strain genotypes used in this study can be found in Table 2.

**Table 2.**
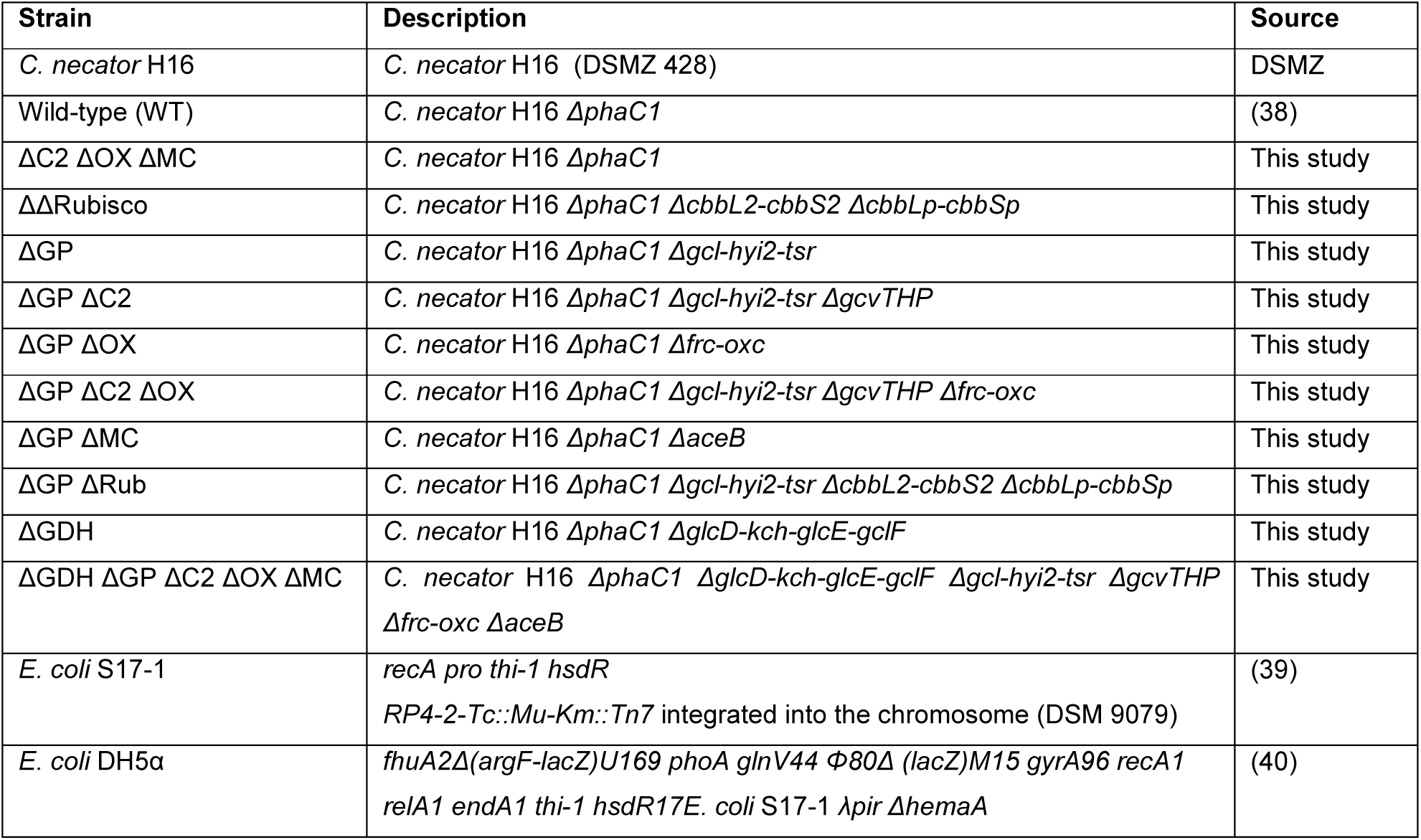
Strains used in this study

| Strain | Description | Source |
| --- | --- | --- |
| *C. necator* H16 | *C. necator* H16 (DSMZ 428) | DSMZ |
| Wild-type (WT) | *C. necator* H16 $\Delta phaC1$ | (38) |
| $\Delta C2 \Delta OX \Delta MC$ | *C. necator* H16 $\Delta phaC1$ | This study |
| $\Delta \Delta Rubisco$ | *C. necator* H16 $\Delta phaC1 \Delta cbbL2-cbbS2 \Delta cbbLp-cbbSp$ | This study |
| $\Delta GP$ | *C. necator* H16 $\Delta phaC1 \Delta gcl-hyi2-tsrl$ | This study |
| $\Delta GP \Delta C2$ | *C. necator* H16 $\Delta phaC1 \Delta gcl-hyi2-tsrl \Delta gcvTHP$ | This study |
| $\Delta GP \Delta OX$ | *C. necator* H16 $\Delta phaC1 \Delta frc-oxc$ | This study |
| $\Delta GP \Delta C2 \Delta OX$ | *C. necator* H16 $\Delta phaC1 \Delta gcl-hyi2-tsrl \Delta gcvTHP \Delta frc-oxc$ | This study |
| $\Delta GP \Delta MC$ | *C. necator* H16 $\Delta phaC1 \Delta aceB$ | This study |
| $\Delta GP \Delta Rub$ | *C. necator* H16 $\Delta phaC1 \Delta gcl-hyi2-tsrl \Delta cbbL2-cbbS2 \Delta cbbLp-cbbSp$ | This study |
| $\Delta GDH$ | *C. necator* H16 $\Delta phaC1 \Delta glcD-kch-glcE-gclF$ | This study |
| $\Delta GDH \Delta GP \Delta C2 \Delta OX \Delta MC$ | *C. necator* H16 $\Delta phaC1 \Delta glcD-kch-glcE-gclF \Delta gcl-hyi2-tsrl \Delta gcvTHP \Delta frc-oxc \Delta aceB$ | This study |
| *E. coli* S17-1 | *recA pro thi-1 hsdR*<br>*RP4-2-Tc::Mu-Km::Tn7* integrated into the chromosome (DSM 9079) | (39) |
| *E. coli* DH5α | *fhuA2\Delta(argF-lacZ)U169 phoA glnV44 \Phi80\Delta(lacZ)M15 gyrA96 recA1 relA1 endA1 thi-1 hsdR17E. coli* S17-1 $\lambda pir \Delta hemaA$ | (40) |

Gene deletions were performed with the pLO3 suicide vector, as previously described (41, 42). Briefly, ~1 kb homology arms upstream and downstream of the gene or operon to be deleted site were PCR amplified (Phusion HF polymerase with DMSO, Thermo Scientific) from genomic *C. necator* DNA using primers as listed in Supplementary Table S1. Homology arms were cloned into pLO3 backbone (SacI, XbaI digested) via In-Fusion Assembly (Takara) and confirmed by Sanger Sequencing (LGC). *C. necator* was conjugated with *E. coli* S17-1 harboring pLO3 knockout vectors, and single-homologous recombination clones were selected on agar plates with tetracycline (10 µg/mL) and 10 µg/mL gentamycin tot counter-select for *E. coli*. Next, some transconjugants were grown in an overnight liquid culture (without tetracycline) to support a second homologous recombination event. The overnight cultures were plated on LB with 10% sucrose to allow for SacB counter selection. Resulting colonies were screened by colony PCR (OneTaq, Thermo Scientific) to identify gene-deleted strains (primers in Supplementary Table S1), which were further verified for having lost tetracycline resistance. Biological duplicates of each knockout strain were constructed.

### Growth medium and conditions

*C. necator* and *E. coli* were cultivated for routine cultivation and genetic modifications on Lysogeny Broth (LB) (1% NaCl, 0.5% yeast extract and 1% tryptone). Routine cultivation was performed in 3 mL medium in 12 mL glass culture tubes in a Kuhner shaker incubator (240 rpm) at 30ºC for *C. necator* and 37ºC for *E. coli.* Growth characterization and transcriptomic experiments of *C. necator* were performed in J Minimal Medium (JMM) as reported previously (43).

### 96-well plates growth experiments

Growth on glycolate or oxalate was performed on JMM with 40 mM sodium glycolate or 40 mM sodium oxalate (Sigma-Aldrich). Experiments were performed in 96 well-plates (Nunc transparent flat-bottom, Thermo Scientific) under ambient air conditions. Precultures for these experiment were performed in JMM with 20 mM fructose in glass tubes, and next washed three times and inoculated at an OD_600_ of 0.01 or 0.005. 150 µL of culture medium was topped with 50 µL of transparent mineral oil (Sigma-Aldrich) to prevent evaporation (O_2_ and CO_2_ can diffuse well through this oil). Plates were incubated with continuous shaking, alternating between 1 minute orbital and 1 minute linear, in a BioTek Epoch 2 plate reader. OD_600_ values were recorded every ~12 minutes. Growth data were processed by an in-house Matlab script, converting OD_600_ measured by the reader to cuvette OD_600_, by multiplication with 4.35. All growth experiments were performed at least in triplicates, and the growth curves shown are representative curves.

### Bioreactor experiments

Autotrophic growth was performed in bioreactor experiments. Strains were precultured in 100 mL JMM in 500 mL erlenmeyers with 80 mM sodium formate and 10% CO_2_ in the headspace. Precultures were washed three times and resuspended in JMM without carbon source. Bioreactors (DASGIP, Eppendorf) with 700 mL JMM medium were inoculated at a starting OD_600_ of 0.05. The reactors were continuously sparged (6.25 L/minute) with a gas mixture of 4% hydrogen and 96% ambient air (or 86% air + 10% CO_2_ for control experiments), controlled by a gas flow controller (HovaGas). The reactors were stirred at 200 rpm and temperature controlled at 30ºC. To compensate for high evaporation losses due to high sparging flow, a level sensor controlled a feed pump of sterile water. Samples for OD_600_ measurements and supernatant analysis were taken daily, while fast-growth at 10% CO_2_ was monitored by an on-line sensor calibrated for cuvette OD_600_.

### RNA isolation and sequencing

For transcriptome analysis, cells were grown in 10 mL JMM (without carbon source) in 100 mL erlenmeyers within a 10 L desiccator filled with 4% hydrogen + 96% ambient air, or 4% hydrogen + 10% CO_2_ and 86% air. To maintain hydrogen and CO_2_, the gas phase in the desiccator was exchanged at least twice a day. Biological replicate cultures were harvested in log phase (2 mL culture for OD_600_ ~0.2 for ambient CO_2_, 1 mL for OD_600_ ~0.4 for 10% CO_2_) and stabilized by RNA Protect Bacteria Kit (Qiagen). Next, cells were lysed using lysozyme and a bead-beating step with glass beads (Retschmill, MM200), for 5 minutes at 30 hertz. RNA was purified using the RNeasy Mini kit (Qiagen) according to manufacturer’s instructions and on-column DNAase digestion (DNase kit Qiagen). rRNA depletion (RiboZero kit), cDNA library preparation, and paired-end 150 bp read sequencing (Illumina HiSeq 3000) was performed by the Max Planck Genome Centre Cologne, Germany.

### Transcriptome data analysis

Sequence data of all samples were mapped with STAR v2.5.4b using default parameters (44). Ensembl version 38 genome reference in FASTA format and Ensembl version 38 cDNA Annotation in GTF format were used for genome indexing with adapted parameters for genome size (--genomeSAindexNbases 10) and read length (--sjdbOverhang 150). Anti-strand reads out of the ReadsPerGene files, which are automatically generated by STAR, were used in two different ways: calculating reads per kilo base of exon per million mapped reads (RPKM) for sample-wise transcript abundances as well as merging in order to perform a differential expression analysis with DESeq2 (45) as guided by the rnaseqGene Bioconductor workflow (https://bioconductor.org/packages/release/workflows/vignettes/rnaseqGene/inst/doc/rnaseqGene.html). Briefly, samples were grouped by single parameter condition (ambient CO_2_ and 10% CO_2_), read count data were then loaded with DESeqDataSetFromMatrix to create a DeSeqDataSet object to subsequently run the standard analysis consisting of the functions DESeq and Results. Then, log2-transformed fold changes (log2fc) for ambient CO_2_ compared to 10% CO_2_ and absolute log10 of adjusted p-values were determined and were visualized in a Volcano plot.

### Supernatant analysis for glycolate and glyoxylate

Glycolate concentrations in culture supernatant were determined by ion chromatography (IC) analysis. Supernatant samples were diluted 1:100 in ultrapure water (Milli-Q). The samples were analyzed in an ICS 3000 (Dionex) ion chromatography system, which was combined with a AS50 auto sampler. The samples were run through a Dionex™ IonPac™ AS11 IC column (4 mm diameter, 250 mm length (044076)) and a guard column (4mm diameter, 50mm length (044078)). Samples were run following KOH eluent gradient: 1 mM from 0 to 5 minutes, 1 mM to 15 mM from 5 to 14 minutes,15 mM to 30 mM from 14 to 23 minutes,30 mM to 60 mM from 23 to 31 min at a flow rate of 0.015 mL/minute. The experimental data were analyzed using Chromeleon 6.8. Concentrations were calculated based on a standard curve generated for sodium glycolate (Sigma-Aldrich) in JMM medium. Glyoxylate concentrations were determined by a colorimetric assay based on a reported protocol (46). Specifically, 216 µL from the supernatant sample were mixed with 24 µL of 1% w/v phenylhydrazine in 0.1 M HCl and incubated for 10 minutes at 60ºC and cooled down. To 100 µL of this mixture, 50 µL concentrated HCl and 20 µL 1.6% w/v potassium ferricyanide were added, while background control samples were prepared with 100 µL of reacted sample mixture with 50 µL concentrated HCl and 20 µL MQ water. These samples were incubated for exactly 12 minutes and then absorbance of 1,5-diphenylformazan at 520 nm was recorded by in a BioTek Epoch 2. Differences in absorbance were calculated for each sample by subtracting absorbance from background controls and glyoxylate concentration could be determined based on a standard curve. All supernatant samples of all autotrophic cultures in this work resulted in negligible levels of <0.1 mM glyoxylate.

### Genomic prevalence analysis of Rubisco and malate synthase

Lists of bacterial genomes containing the large subunit of Rubisco (PF00016) or malate synthase (PF01274) were downloaded from the AnnoTree website (47) on April 27^th^h 2020 by searching for the appropriate protein families. As of writing, AnnoTree uses version 89 of the GTDB taxonomy (48), which was retrieved from the GTDB website on the same date. Rubisco sequences were filtered by using usearch (49) to remove any amino acid sequences with >30% identity to known Rubisco-like proteins (RLPs, type IV Rubiscos). The list of Rubisco-like proteins was drawn from (50) and Rubisco-like proteins were removed because they do not catalyze the carboxylation reaction (51). Organisms encoding R and malate synthase were identified by using their GTDB IDs to merge the two tables along with taxonomic information. This permitted calculation of the fraction of Rubisco-containing genomes that also contain malate synthase for each order in the GTDB taxonomy. Analyses were performed in a custom Python script that is available here: https://github.com/flamholz/malate_synthase/blob/master/pipeline/01_plot_co_occurrence.ipynb.

## Acknowledgements

We thank Patrick Behrens, Viswanada Bysani and Natalia Giner Laguarda for assistance with the bioreactor experiments and sampling. We thank Leonardo Perez de Souza, Alisdair Fernie, and Elad Noor for fruitful discussions and advice on this work. We also thank Ari Satanowksi, Armin Kubis and Sebastian Wenk and for feedback on the manuscript and helpful suggestions. We thank the Max Planck Genome Centre Cologne (http://mpgc.mpipz.mpg.de/home/) for performing the RNA-sequencing in this study. This project was funded by the Max Planck Society. N.J.C. is funded by the Dutch Organization of Science (NWO) by a Rubicon (019.163LW.035) and a Veni (VI.Veni.192.156) fellowship. S.F. and O.L. thank for funding by the Deutsche Forschungsgemeinschaft (DFG, German Research Foundation) under Germany’s Excellence Strategy – EXC 2008 – 390540038 – UniSysCat.

## Author contributions

This study was conceived by N.J.C. and A.B.-E. Experiments were designed by N.J.C., G.S., A.F., A.I.F., S.F., O.L. and A.B.-E. Experiments were performed by N.J.C., G.S., W.N. and S.F. Data analysis was performed by N.J.C., G.S.,A.F. and A.I.F. The manuscript was written by N.J.C. and A.B.-E. All authors read and approved the manuscript.

## Data availability

Transcriptomic data are available in the Gene Expression Omnibus (GEO, https://www.ncbi.nlm.nih.gov/geo/) under accession GSE141999 and are available in Supplementary Data 1. Bioinformatic genome analysis of Rubisco and malate synthase are available in Supplementary Data 2 and https://github.com/flamholz/malate_synthase/tree/master/pipeline.

## References

1. Flügel F, Timm S, Arrivault S, Florian A, Stitt M, Fernie AR. 2017. The Photorespiratory Metabolite 2-Phosphoglycolate Regulates Photosynthesis and Starch Accumulation in Arabidopsis. Plant Cell 29:2537–2551.

2. Anderson LE. 1971. Chloroplast and cytoplasmic enzymes II. Pea leaf triose phosphate. Biochim Biophys Acta 235:237–244.

3. Bauwe H, Hagemann M, Fernie AR. 2010. Photorespiration: players, partners and origin. Trends Plant Sci 15:330–336.

4. Fernie AR, Bauwe H, Eisenhut M, Florian A, Hanson DT, Hagemann M, Keech O, Weber APM, Westhoff P. 2013. Perspectives on plant photorespiratory metabolism. Plant Biol 15:748–753.

5. Eisenhut M, Ruth W, Haimovich M, Bauwe H, Kaplan A, Hagemann M. 2008. The photorespiratory glycolate metabolism is essential for cyanobacteria and might have been conveyed endosymbiontically to plants. Proc Natl Acad Sci U S A 105:17199–17204.

6. Nakamura Y, Kanakagiri S, Van K, He W, Spalding MH. 2005. Disruption of the glycolate dehydrogenase gene in the high-CO_2_-requiring mutant HCR89 of *Chlamydomonas reinhardtii*. Can J Bot 83:820–833.

7. Kern R, Bauwe H, Hagemann M. 2011. Evolution of enzymes involved in the photorespiratory 2-phosphoglycolate cycle from cyanobacteria via algae toward plants. Photosynth Res 109:103–114.

8. Eisenhut M, Kahlon S, Hasse D, Ewald R, Judy Lieman-Hurwitz TO, Ruth W, Bauwe H, Kaplan A, Hagemann M. 2006. The Plant-Like C2 Glycolate Cycle and the Bacterial-Like Glycerate Pathway Cooperate in Phosphoglycolate Metabolism in Cyanobacteria. Plant Physiol 142:333–342.

9. Forrester ML, Krotkov G, Nelson CD. 1966. Effect of Oxygen on Photosynthesis, Photorespiration and Respiration in Detached Leaves. I. Soybean. Plant Physiol 41:422–427.

10. Raven JA. 2009. Contributions of anoxygenic and oxygenic phototrophy and chemolithotrophy to carbon and oxygen fluxes in aquatic environments. Aquat Microb Ecol 56:177–192.

11. Claassens NJ, Sánchez-Andrea I, Sousa DZ, Bar-Even A. 2018. Towards sustainable feedstocks: A guide to electron donors for microbial carbon fixation. Curr Opin Biotechnol 50:195–205.

12. Bowien B, Kusian B. 2002. Genetics and control of CO_2_ assimilation in the chemoautotroph *Ralstonia eutropha*. Arch Microbiol 178:85–93.

13. Gai CS, Lu J, Brigham CJ, Bernardi AC, Sinskey AJ. 2014. Insights into bacterial CO_2_ metabolism revealed by the characterization of four carbonic anhydrases in *Ralstonia eutropha* H16. AMB Express 4:2.

14. Cramm R. 2009. Genomic view of energy metabolism in *Ralstonia eutropha* H16. J Mol Microbiol Biotechnol 16:38–52.

15. Badger MR, Bek EJ. 2008. Multiple Rubisco forms in proteobacteria: Their functional significance in relation to CO_2_ acquisition by the CBB cycle. J Exp Bot 59:1525–1541.

16. Satagopan S, Tabita FR. 2016. RubisCO selection using the vigorously aerobic and metabolically versatile bacterium *Ralstonia eutropha*. FEBS J 283:2869–2880.

17. Flamholz AI, Prywes N, Moran U, Davidi D, Bar-On YM, Oltrogge LM, Alves R, Savage D, Milo R. 2019. Revisiting Trade-offs between Rubisco Kinetic Parameters. Biochemistry 58:3365–3376.

18. Codd GA, Bowien B, Schlegel HG. 1976. Glycollate production and excretion by *Alcaligenes eutrophus*. Arch Microbiol 110:167–171.

19. King WR, Andersen K. 1980. Efficiency of CO2 Fixation in a Glycollate Oxidoreductase Mutant of Alcaligenes eutrophus which Exports Fixed Carbon as Glycollate. Arch Microbiol 90:84–90.

20. Schaferjohann J, Yoo JG, Kusian B, Bowien B. 1993. The *cbb* operons of the facultative chemoautotroph Alcaligenes eutrophus encode phosphoglycolate phosphatase. J Bacteriol 175:7329–7340.

21. Pohlmann A, Fricke WF, Reinecke F, Kusian B, Liesegang H, Cramm R, Eitinger T, Ewering C, Pötter M, Schwartz E, Strittmatter A, Voss I, Gottschalk G, Steinbüchel A, Friedrich B, Bowien B. 2006. Genome sequence of the bioplastic-producing “Knallgas” bacterium Ralstonia eutropha H16. Nat Biotechnol 24:1257–62.

22. Schneider K, Skovran E, Vorholt JA. 2012. Oxalyl-coenzyme A reduction to glyoxylate is the preferred route of oxalate assimilation in *Methylobacterium extorquens AM1*. J Bacteriol 194:3144–3155.

23. Borzyskowski LS Von, Severi F, Krüger K, Hermann L, Gilardet A, Sippel F, Pommerenke B, Claus P, Cortina NS, Glatter T, Zauner S, Zarzycki J, Fuchs BM, Bremer E, Maier UG, Amann RI, Erb TJ. 2019. Marine Proteobacteria metabolize glycolate via the β-hydroxyaspartate cycle. Nature 575:500–504.

24. Brigham CJ, Budde CF, Holder JW, Zeng Q, Mahan AE, Rha CK, Sinskey AJ. 2010. Elucidation of β-oxidation pathways in *Ralstonia eutropha* H16 by examination of global gene expression. J Bacteriol 192:5454–5464.

25. Wang ZX, Brämer CO, Steinbüchel A, Bra CO, Steinbu A, Brämer CO, Steinbüchel A. 2003. The glyoxylate bypass of *Ralstonia eutropha*. FEMS Microbiol Lett 228:63–71.

26. Howard BR, Endrizzi JA, Remington SJ. 2000. Crystal structure of *Escherichia coli* malate synthase G complexed with magnesium and glyoxylate at 2.0 Å resolution: Mechanistic implications. Biochemistry 39:3156–3168.

27. Flamholz A, Noor E, Bar-Even A, Milo R. 2012. EQuilibrator - The biochemical thermodynamics calculator. Nucleic Acids Res 40:D770–775.

28. Erb TJ, Zarzycki J. 2016. Biochemical and synthetic biology approaches to improve photosynthetic CO_2_-fixation. Curr Opin Chem Biol 34:72–79.

29. South PF, Cavanagh AP, Liu HW, Ort DR. 2019. Synthetic glycolate metabolism pathways stimulate crop growth and productivity in the field. Science (80-) 363:eaat9077.

30. Maier A, Fahnenstich H, von Caemmerer S, Engqvist MKM, Weber APM, Flügge U-I, Maurino VG. 2012. Transgenic Introduction of a Glycolate Oxidative Cycle into *A. thaliana* Chloroplasts Leads to Growth Improvement. Front Plant Sci 3:1–12.

31. Kubis A, Bar-even A. 2019. Synthetic biology approaches for improving photosynthesis. J Exp Bot 70:1425–1433.

32. Knoop H, Grudel M, Zilliges Y, Lehmann R, Hoffmann S, Lockau W, Steuer R. 2013. Flux Balance Analysis of Cyanobacterial Metabolism: The Metabolic Network of *Synechocystis* sp. PCC 6803. PLoS Comput Biol 9:e1003081.

33. Le Y, Berla B, He L, Pakrasi HB, Tang YJ. 2014. 13C-MFA delineates the photomixotrophic metabolism of *Synechocystis* sp. PCC 6803 under light- and carbon-sufficient conditions. Biotechnol J 9:684–692.

34. Gründel M, Knoop H, Steuer R. 2017. Activity and functional properties of the isocitrate lyase in the cyanobacterium *Cyanothece* sp. PCC 7424. Microbiology 163:731–744.

35. Zhang S, Bryant DA. 2015. Biochemical validation of the glyoxylate cycle in the cyanobacterium *Chlorogloeopsis fritschii* strain PCC 9212. J Biol Chem 290:14019–14030.

36. Norman EG, Colman B. 1992. Formation and metabolism of glycolate in the cyanobacterium *Coccochloris peniocystis*. Arch Microbiol 157:375–380.

37. Eley JH. 1988. Glyoxylate cycle enzyme activities in the cyanobaceterium *Anacystis Nidulans*. J Phycol 24:586–588.

38. Lütte S, Pohlmann A, Zaychikov E, Schwartz E, Becher JR, Heumann H, Friedrich BBBB, Lutte S, Pohlmann A, Zaychikov E, Schwartz E, Becher JR, Heumann H, Friedrich BBBB, Lütte S, Pohlmann A, Zaychikov E, Schwartz E, Becher JR, Heumann H, Friedrich BBBB. 2012. Autotrophic production of stable-isotope-labeled arginine in *Ralstonia eutropha* strain H16. Appl Environ Microbiol 78:7884–7890.

39. Simon R, Priefer U, Puhl A. 1983. A broad host range mobilization system for in vivo genetic engineering: transposon mutagenesis in gram negative bacteria. Nat Biotechnol 784–791.

40. Thoma S, Schobert M. 2009. An improved *Escherichia coli* donor strain for diparental mating. FEMS Microbiol Lett 294:127–132.

41. Lenz O, Friedrich B. 1998. A novel multicomponent regulatory system mediates H_2_ sensing in *Alcaligenes eutrophus*. Proc Natl Acad Sci USA 95:12474–12479.

42. Lenz O, Schwartz E, Dernedde J. 1994. The *Alcaligenes eutrophus* H16 *hoxX* gene participates in hydrogenase regulation. J Bacteriol 176:4385–4393.

43. Li H, Opgenorth PH, Wernick DG, Rogers S, Wu T-Y, Higashide W, Malati P, Huo Y-X, Cho KM, Liao JC. 2012. Integrated Electromicrobial Conversion of CO_2_ to Higher Alcohols. Science 335:1596–1596.

44. Dobin A, Davis CA, Schlesinger F, Drenkow J, Zaleski C, Jha S, Batut P, Chaisson M, Gingeras TR. 2013. STAR: ultrafast universal RNA-seq aligner. Bioinformatics 29:15–21.

45. Love MI, Huber W, Anders S. 2014. Moderated estimation of fold change and dispersion for RNA-seq data with DESeq2. Genome Biol 15:1–21.

46. Rojano-Delgado AM, Priego-Capote F, Castro MDL De, Prado R De. 2010. Screening and confirmatory analysis of glyoxylate: A biomarker of plants resistance against herbicides. Talanta 82:1757–1762.

47. Mendler K, Chen H, Parks DH, Lobb B, Hug LA, Doxey AC. 2019. Annotree: Visualization and exploration of a functionally annotated microbial tree of life. Nucleic Acids Res 47:4442–4448.

48. Parks DH, Chuvochina M, Waite DW, Rinke C, Skarshewski A, Chaumeil PA, Hugenholtz P. 2018. A standardized bacterial taxonomy based on genome phylogeny substantially revises the tree of life. Nat Biotechnol 36:996.

49. Edgar RC. 2010. Search and clustering orders of magnitude faster than BLAST. Bioinformatics 26:2460–2461.

50. Jaffe AL, Castelle CJ, Dupont CL, Banfield JF. 2019. Lateral Gene Transfer Shapes the Distribution of RuBisCO among Candidate Phyla Radiation Bacteria and DPANN Archaea. Mol Biol Evol 36:435–446.

51. Tabita FR, Satagopan S, Hanson TE, Kreel NE, Scott SS. 2008. Distinct form I, II, III, and IV Rubisco proteins from the three kingdoms of life provide clues about Rubisco evolution and structure/function relationships. J Exp Bot 59:1515–1524.

