## Supplementary Fig S1 for "Photorespiration pathways in a chemolithoautotroph"

Supplementary figure 1

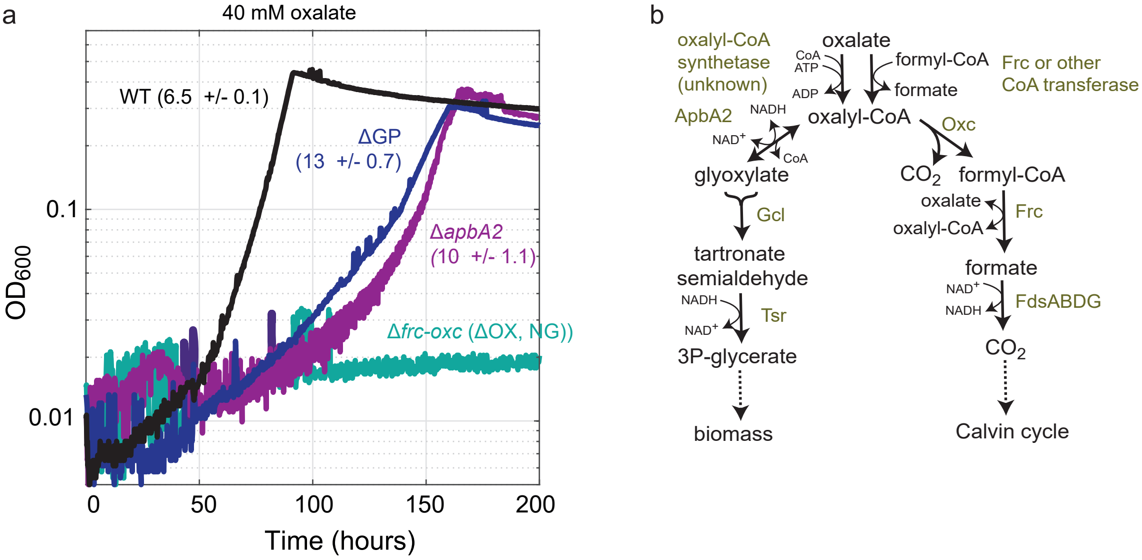


**Supplementary Figure S1. Growth of *C. necator* on oxalate. (a)** Growth experiment of gene-deletion strains on oxalate**.** Growth experiments were conducted in 96-well plate readers in minimal medium (JMM) supplemented with 40 mM sodium oxalate. Doubling times (hours) and standard deviations of triplicates are shown in between brackets, 'NG' corresponds to 'no growth'. Growth experiments were performed in biological triplicates and showed identical growth curves (±5%); hence, representative curves are shown. **(b)** Growth on oxalate can proceed via two main routes in *C. necator*, the reduction of oxalyl-CoA to glyoxylate and its assimilation via the glycerate route, or via the oxidation of oxalyl-CoA and full decarboxylation to $CO_2$. Growth of gene-deleted strains for one of both routes show that both play a role in growth on oxalate, the decarboxylative route seems essential as its knockout abolished growth. Abbreviations: ApbA2, CoA-acylating glyoxylate dehydrogenase; FdsABDG, formate dehydrogenase complex; Frc, formyl-CoA transferase; Gcl, glyoxylate carboligase; ΔGP, glycerate pathway knockout (*Δgcl-hyi-tsr*); ΔOX, oxalate decarboxylation knockout (*frc-oxc*); Oxc, oxalyl-CoA decarboxylase; Tsr, tartronate semialdehyde reductase; TtuD1, glycerate kinase; and WT, wild-type.
