## Supplementary Table S1 for "Photorespiration pathways in a chemolithoautotroph"

**Supplementary Table S1. Primers used in this study.**

| **Gene/operon deletion** | **Gene ID(s)** | **Primer type** | **Primer sequence 5’ – 3’** |
| --- | --- | --- | --- |
| *ΔgclD-kch-EF*  (ΔGDH) | H16_A3094-3097 | Fw_HRup | cacctagatccttttaattcctggaggtggaatcggtct |
|  |  | Rv_HRup | tgtcgaccagattcatgaacgactcctgtg |
|  |  | Fw_HRdown | ccacctccagctggtcgacaagatgctcggcta |
|  |  | Rv_HRdown | gtttaaacagtcgactctagttggtttgccatgttgatgt |
|  |  | Fw_colPCR_1 | gaggacggcactgacaagac |
|  |  | Fw_colPCR_2 | gacaagctcgacatggtctg |
|  |  | Rv_colPCR | atcatgcctgctctcctttc |
| *Δgcl-hyi2-tsr*  (ΔGP) | H16_A3598-3600 | Fw_HRup | acctagatccttttaattcgtccatcggatattcgtctcc |
|  |  | Rv_HRup | ggtggttagagctctcatctttgccatgatct |
|  |  | Fw_HRdown | agatgagagctctaaccaccagatcgccaag |
|  |  | Rv_HRdown | gtttaaacagtcgactctagattacgcagggcttgctg |
|  |  | Fw_colPCR | actccaacggaattcacacc |
|  |  | Rv_colPCR_1 | cagcaggtcgatatgctcag |
|  |  | Rv_colPCR_2 | aaatcaggcacagcaccag |
| *ΔgcvT1HP*  (ΔC2) | H16_A3619-3621 | Fw_HRup | cacctagatccttttaattcggcaagggccgtccaggtag |
|  |  | Rv_HRup | cgtagtcgctcatggaatcctctggggcagtc |
|  |  | Fw_HRdown | ggattccatgagcgactacgtggtggactgag |
|  |  | Rv_HRdown | gtttaaacagtcgactctaggttctcttcattgactgcgatgg |
|  |  | Fw_colPCR | ggtatcgaggacagaggggtca |
|  |  | Rv_colPCR | caggtcagctcgttttccat |
| *frc-oxc* (ΔOX) | H16_B1711-1712 | Fw_HRup | acctagatccttttaattcgatagagctggcccaccgtagc |
|  |  | Rv_HRup | agtcggaaacgagtcccaggtgatctgacgc |
|  |  | Fw_HRdown | cctgggactcgtttccgacttctgccatgg |
|  |  | Rv_HRdown | gtttaaacagtcgactctagttcctgtcggacaagttcgg |
|  |  | Fw_colPCR | tctggtctgtcatggggatt |
|  |  | Rv_colPCR | acttgaagggctaggggaaa |
| *ΔaceB*  (ΔMC) | H16_A2217 | Fw_HRup | acctagatccttttaattcgcgattaagcccgtaggcgacc |
|  |  | Rv_HRup | aagactgatgctgacgctgccgctgtatgag |
|  |  | Fw_HRdown | gcagcgtcagcatcagtcttctcctgtgatc |
|  |  | Rv_HRdown | gtttaaacagtcgactctaggtctccagttgtttgaagtgg |
|  |  | Fw_colPCR | ccctacccgcttaaacttcc |
|  |  | Rv_colPCR | caggccctcatgaacaagat |
| *ΔcbbLS2*  (ΔRub) | H16_B1394-1395 | Fw_HRup | acctagatccttttaattcgatggtggtcagggtatggtc |
|  |  | Rv_HRup | ccaagccaagcatgcgctactcgatcgaga |
|  |  | Fw_HRdown | gtagcgcatgcttggcttggaccgattcag |
|  |  | Rv_HRdown | gtttaaacagtcgactctagacatatgcgcaacatgccagatg |
|  |  | Fw_colPCR | atcaatcgagcatccgactc |
|  |  | Rv_colPCR | acccctttgcatcgtgtaac |
| *ΔcbbLSp*  (ΔRub) | PHG_426-427 | Fw_HRup | acctagatccttttaattcgtttcgtcgtcaaagcggtac |
|  |  | Rv_HRup | gacaagcatgtattcgatcgagagctacgcc |
|  |  | Fw_HRdown | cgatcgaatacatgcttgtctccttgcgtg |
|  |  | Rv_HRdown | gtttaaacagtcgactctagaatcccctaggccacaagcc |
|  |  | Fw_colPCR | ggcttcagtccgatcagttc |
|  |  | Rv_colPCR | aggaaggacatgagcagtgg |
| *ΔapbA2* | H16_B1719 | Fw_HRup | acctagatccttttaattcgcaaaacggcggaccaaggtc |
|  |  | Rv_HRup | gacagcaatgctgatcaagcagcgcgacac |
|  |  | Fw_HRdown | gcttgatcagcattgctgtctccttgatgg |
|  |  | Rv_HRdown | gtttaaacagtcgactctagagaactgtaagagacgccgc |
